# Inferring Reaction Networks using Perturbation Data

**DOI:** 10.1101/351767

**Authors:** Kiri Choi, Joesph Hellerstein, H. Steven Wiley, Herbert M. Sauro

**Affiliations:** Department of Bioengineering, William H. Foege Building, Box 355061, Seattle, WA, USA; eScience Institute, University of Washington, Seattle, Washington, USA; Environmental Molecular Sciences Laboratory, Pacific Northwest National Laboratory, Richland, WA, USA

## Abstract

In this paper we examine the use of perturbation data to infer the underlying mechanistic dynamic model. The approach uses an evolutionary strategy to evolve networks based on a fitness criterion that measures the difference between the experimentally determined set of perturbation data and proposed mechanistic models. At present we only deal with reaction networks that use mass-action kinetics employing uni-uni, bi-uni, uni-bi and bi-bi reactions. The key to our approach is to split the algorithm into two phases. The first phase focuses on evolving network topologies that are consistent with the perturbation data followed by a second phase that evolves the parameter values. This results in almost an exact match between the evolved network and the original network from which the perturbation data was generated from. We test the approach on four models that include linear chain, feed-forward loop, cyclic pathway and a branched pathway. Currently the algorithm is implemented using Python and libRoadRunner but could at a later date be rewritten in a compiled language to improve performance. Future studies will focus on the impact of noise in the perturbation data on convergence and variability in the evolved parameter values and topologies. In addition we will investigate the effect of nonlinear rate laws on generating unique solutions.

## Introduction

In [8, 2] we illustrated how it was possible to use an evolutionary approach to generate biochemical networks that had specific dynamic behaviors. This was done by defining an objective function and then use a genetic algorithm to evolve networks that possessed the desired behavior. For example, we demonstrated how to evolve networks that showed sustained oscillations, could compute square and cube roots, showed bi-stability, and behaved as low, high and bandpass filters.

This paper extends our previous work by using an evolutionary approach to infer a network’s structure and kinetics using perturbation data. Instead of evolving networks with specific dynamic behaviors we evolve fully mechanistic networks that are consistent with a given set of perturbation data. Modular response analysis (MRA) has used perturbation data to infer networks and dynamics [7, 3]. A recent review describing the development of this technique can be found in [9]. This method relies on using perturbation data to reconstruct the Jacobian matrix from which the modular structure of the network can be inferred. The approach is promising but it can be difficult to use it to reconstruct a detailed mechanistic model. Instead, the approach we propose can be used to infer detailed mechanistic models from perturbation data. Moreover, noise in the perturbation data will lead naturally to the generation of multiple models that are consistent with the data, something MRA cannot easily do. However, the effect of noise will be discussed in a later revision.

Consider a biochemical network with *n* reactions and *m* species at steady state. We define a perturbation of the *m*_*i*_ species as the change in the steady state level of *m*_*i*_ as a result of a change to the reaction *n*_*i*_. We will define perturbations using logarithmic gains similar to those defined in metabolic control analysis [6, 11], also known as control coefficients. Metabolic control analysis (MCA) studies the sensitivity of a chemical reaction network to perturbations and how the perturbations propagate through the network. A control coefficient describes how a flux *J* or a species concentration *S* is affected by changes to the concentration of an enzyme *E*_*i*_. Therefore, flux and concentration control coefficients are often defined using scaled derivative as shown in equations (1).

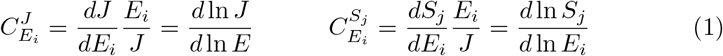

In principle it is possible to experimentally estimate flux and concentration control coefficients for every combination of species and reaction, generating matrices of control coefficients [4].

In this paper, we propose a novel approach to infer dynamic mechanistic models from concentration control coefficients using an evolutionary algorithm. We test the algorithm with four different types of synthetic networks and demonstrate that the algorithm can fully reconstruct the network and its dynamics.

## Methods

The algorithm is iterative in nature but begins by generating a population of random chemical reaction networks with a given number of floating species and reactions. Information on concentration control coefficients is provided to the algorithm in the form of a matrix where the number of rows and columns correspond to the number of species and reactions. For the scope of this paper, we limit our algorithm to inferring networks with Uni-Uni, Uni-Bi, Bi-Uni, and Bi-Bi reactions using mass-action kinetics ^1^. In future work we will consider non-linear Michaelis and Hill like rate laws [10].

The algorithms works iteratively on a population of networks one generation at a time. For each network in the population, the concentration control coefficients as well as the flux and steady-state concentrations of the floating species are determined. The latter two measurements are used to reduce uncertainty in parameter values. The concentration control coefficient matrices are compared with that of the experimentally determined control coefficients of the unknown network. The comparison is done by computing the distance as a Frobenius norm, which is given by equation (2), where *C*^*E*^ is the experimentally determined control coefficient matrix and *C*^*S*^ is the simulated control coefficient matrix. This technique assumes that the number of reactions and species in the unknown network are known. We will consider variable number of reactions in a subsequent revision of the paper.

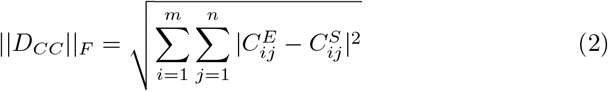

The full distance metric, or fitness, is described in equation (3). Here, SS are the steady-state values and J the steady state flux. Constants *k*_*F*_ and *k*_*S*_ are weights. The subscript *E* refers to the experimental data and *F* to the same data but in the individuals in the population.

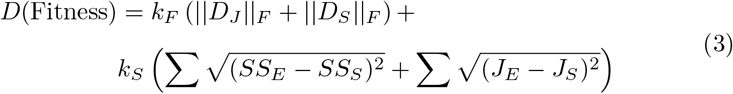

*D* is the matrix of concentration control coefficients, *D*_*J*_ the matrix of fluxes, and *D*_*S*_ the vector of species concentrations.

Moving form one generation to the next involves the following sequence of operations. The fitness of each model is computed using the Frobenius norm (3) [5] and the population of models in the *i*^th^ generation are ranked from best to worst. Assuming that the maximum population size if *N*, then a percentage, *e*_*p*_, of the best individuals in the *i*^th^ generation of *N* individuals is passed to the next generation, *i* + 1^th^. A percentage, *p*_*p*_, of models from the *i*^th^ generation are selected using either roulette, tournament selection or fitness proportionate selection, where the inverse of distance is used as the fitness, and copied to the *i* + 1^th^ generation. The various selection approaches are selectable by the user. Of the *p*_*p*_ models, each is mutated in one of two ways. Mutation can include a change of either a reaction or a rate constant. For the best models in the population we only mutate a reaction. This is to ensure that the mutated output will always be a different structural model compared to one of the best models. Therefore, instead of aiming for the best fitness overall, we aim for best competition between different models, where the goal is to move into the top spot. This effectively minimizes the issue with local minima since our distance metric is affected by both the topology and rate constants of the model. Once the best models and mutated models are passed on to the next generation, the remaining space, *r*_*p*_ percent is filled with freshly generated random networks. Note that the sum *e*_*p*_ + *p*_*p*_ + *r*_*p*_ = 100%. The entire process is repeated for a predetermined number of generations or until the fitness meets a threshold. Figure 1 illustrates the general work flow of the algorithm.

**Figure 1:**
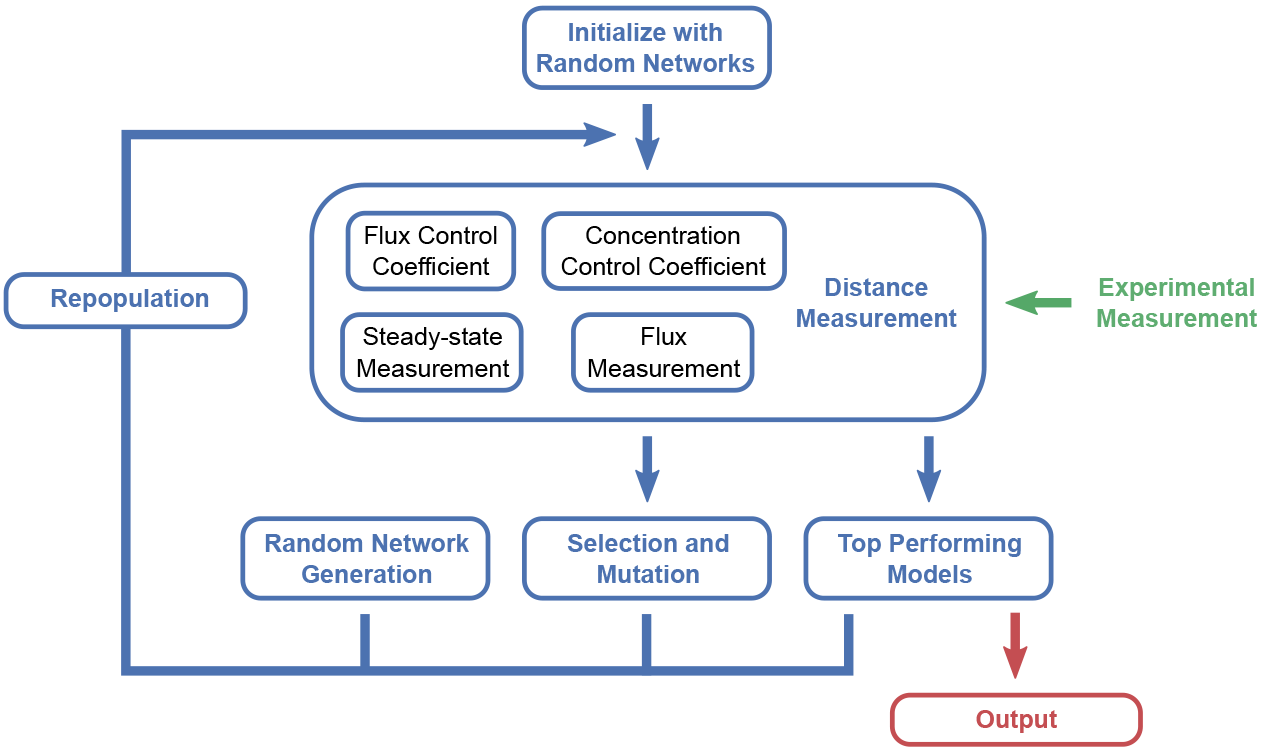
A work-flow diagram illustrating the algorithm. The loop is continued for a set number of generations or until a given fitness threshold is met.

All evolution experiments were developed in Python. For simulation we used libRoadRunner [12]. To improve performance the code could be rewritten in a compiled language such as C/C++. Note that the libRoadRunner is already writen in C/C++.

## Results

**Figure 2:**
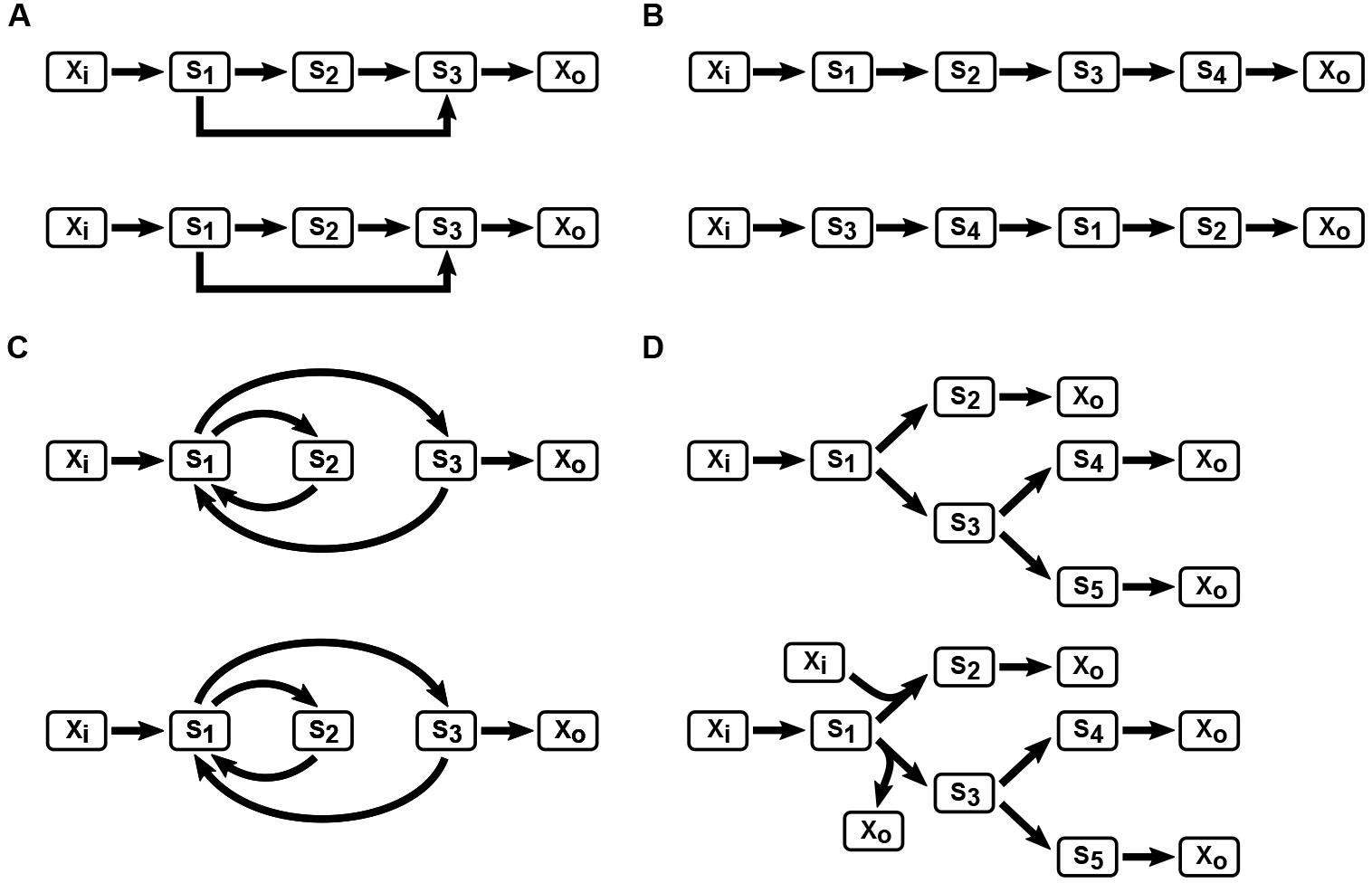
Models used to generate synthetic experimental data and the best performing output acquired from the algorithm using it. Four models are used, including (A) coherent type 1 feed-forward loop, (B) linear chain, (C) nested cycles, and (D) branched pathways. Within each panel, the diagram on the top represents the original model used to generate the synthetic experimental data and diagram below is the model identified by the algorithm as the best performing model. *X*_*i*_ and *X*_*o*_ represent the boundary species which are fixed during a simulation.

To demonstrate the capability of the algorithm, we have tested it on a number of synthetic networks involving a simple feed-forward loop, a linear chain, nested cycles, and multiple branches as illustrated in Figure 2. For evolution runs, the population size was set to 200 networks. Within each generation, the top ten percent of models were passed to the next generation and half of the population selected and mutated to the next generation. This means that 40% of the next generation population was populated with random networks.

Figure 2 shows the top performing models after running the algorithm. For motifs such as feed-forward loop and nested cycles, the algorithm is able to fully recover the original network topology and dynamics. For linear chains, the order of species differs from the original network. This is due to a unique characteristic of linear chains. These alternative models have almost identical flux and concentration control coefficients. For the type of experimental measurements we supply to the algorithm, it is impossible to distinguish these linear chains with differing orders. Addition of time-course measurements does not solve this issue either, and the only potential way to distinguish these networks is to use perturbation studies. Finally, for the branched pathways, the algorithm was able to accurately recover the original model. Of interest are the non-functional components of the evolved network. For example, the second and third reaction in the evolved branched model (D) include the boundary species *X*_1_ and *X*_*o*_ which were not present in the original model. However, these have no effect on the network dynamics since the effect of a particular boundary species can be absorbed into the rate constant for the reaction. Figure 3 shows the change in the fitness observed in the population over a number of generations. The number below and on the right side of the *x* axis shows the approximate time it took to run the algorithm.

**Figure 3:**
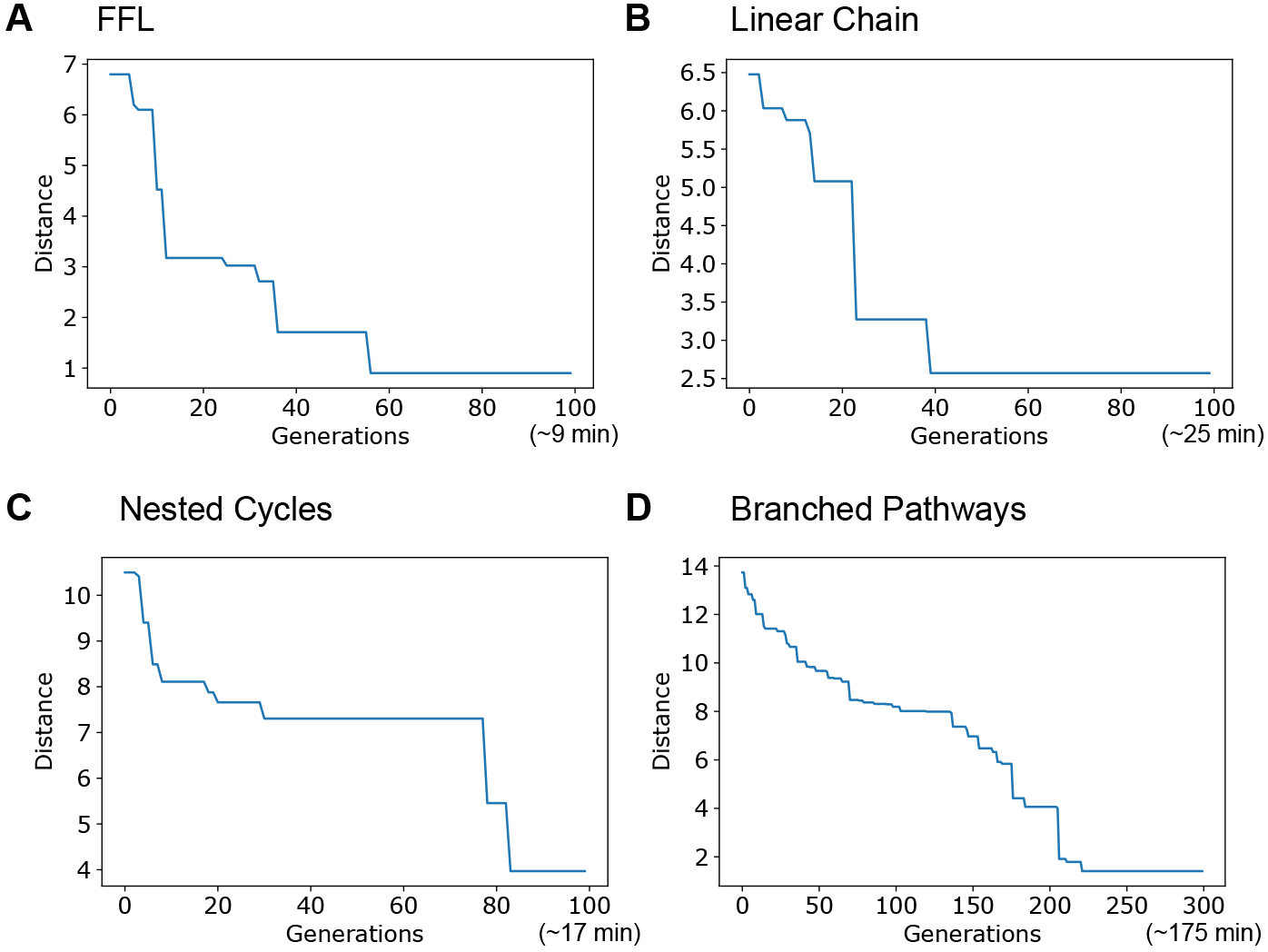
Convergence curves for four models tested in Figure 1. Lines correspond to the fitness of top performing model. Numbers below and to the right of each plot represents the time it took to complete the run.

One point of interest is that in many cases, experimentally measuring flux control coefficients or fluxes are relatively difficult. We report that in our preliminary analysis, using only the Frobenius norm of differences in concentration control coefficient (∥*D*_*S*_∥_*F*_) and steady-state measurement is good enough to recover a large part of the original model, although the time it takes to converge is much longer.

**Figure 4:**
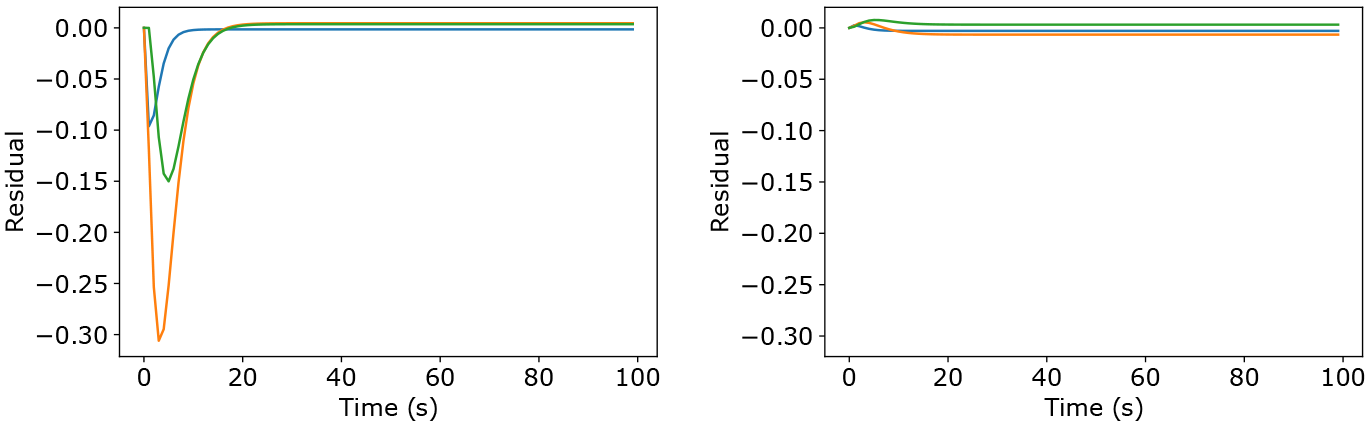
Time-course simulation showing the difference between the best performing model and the original feed-forward loop model. Left is the result of using steady-state measurements only during the second round of iterations. Right is the result of using steady-state measurements and fluxes during the second round of iterations. Blue, orange, and green lines correspond to species S1, S2, and S3, respectively.

We also tested the ability of the evolved models to match the transient behavior of the model. For this, we modified our work-flow slightly, starting with only using control coefficients as the distance metric to recover the stoichiometry, and then running the best model through another round of iteration only using the difference in steady-state measurements. This change separates the entire process into two distinct rounds of iterations, where you first round of iterations look for the correct stoichiometry and then look for the correct parameterization. When only the concentration control coefficients and steady-state measurements are used, the evolved networks were unable to recreate the transient behavior. However, once we included steady-state solutions as well as fluxes in the second round of iterations, the process was able to find the correct set of parameters and the transient behavior was fully recovered, as shown in Figure 4. This may not be unexpected given that the raw laws are linear. In a subsequent analysis we will consider non-linear rate laws. In this situation the transients are unlikley to match and we will consider additional evolutionary curation.

All computation was done on a single core of a AMD Ryzen 1700X machine clocked to 3.4 GHz with 16 GB RAM. All computation was done using libRoadRunner [12] simulator using the Tellurium [1] environment.

## Conclusion

In this paper, we present an effective way of generating an ensemble of reaction network models from perturbation data. We also demonstrate its effectiveness by testing the algorithm on a set of synthetic networks. Although we have only presented the top performing models here, in many cases one will analyze a number of top performing models, potentially using the models as a whole as an ensemble for better predictions. The performance of the algorithm is reasonable, usually converging to a solution on the scale of tens of minutes. For a larger networks, obviously it will take longer to find a good set of solutions, but the entire process is easily parallelizable, which should produce a significant performance boost.

## Availability

Scripts used to generate the results are available at https://github.com/kirichoi/CCR. We recommend installing Tellurium to run the script. Tellurium is available at http://tellurium.analogmachine.org.

## Authors contributions

KC designed the algorithm, generated and analyzed the result, and wrote the code and article; JH read the manuscript and assisted with algorithm development; HSW read the manuscript and conceived the idea of measuring perturbations; HMS conceived the idea and wrote the code and article.

## Acknowledgements

This work was supported by the National Institute of General Medical Sciences of the National Institutes of Health under awards R01-GM081070, R01-GM123032. The content is solely the responsibility of the authors and does not necessarily represent the official views of the National Institutes of Health or National Science Foundation.

## Footnotes

1 Uni-Uni refers to reactions of the type A → B, Uni-Bi refers to reactions of the type A → B + C, Bi-Uni refers to reactions of the type A + B → C, and Bi-Bi refers to reactions of the type A + B → C + D.

